# Adoptive Transfer of ILC2s into the Brain Reveals Tumor Homing in Glioblastoma: A Proof-of-Concept for Cellular Therapy

**DOI:** 10.64898/2025.12.22.695985

**Authors:** Lei P Wang, Bidhan Bhandari, Sahar Emami Naeini, Jack C Yu, Ali S Arbab, Nancy Young, Évila Lopes Salles, Babak Baban

## Abstract

Glioblastoma (GBM) is an aggressive brain tumor with limited treatment options and poor immune cell infiltration. Cellular immunotherapies have transformed cancer treatment but remain largely ineffective against GBM due to the restrictive blood–brain barrier (BBB) and its profoundly immunosuppressive microenvironment. Innate lymphoid cells type 2 (ILC2s) have recently emerged as potential candidates for cellular immunotherapy because of their regenerative and immunomodulatory functions. Here, we provide the first in vivo evidence that systemically administered ILC2s can cross the BBB and home to glioblastoma. Bone marrow–derived ILC2s from C57BL/6 mice were labeled with CFSE and intravenously transferred into hosts bearing orthotopic, luciferase-expressing GL261 tumors. Fluorescence imaging and flow cytometry analyses indicated that transferred ILC2s successfully entered the brain, localized within tumor tissue, and were also detected in meninges and peripheral organs. Although no measurable reduction in tumor growth was observed, these findings support a proof-of-concept for adoptive ILC2 transfer as a feasible approach for targeting CNS tumors and exploring their immunomodulatory potential in GBM.

## Introduction

Glioblastoma The Glioblastoma (GBM) remains one of the most aggressive primary brain tumors, characterized by rapid progression, therapeutic resistance, and a profoundly immuno-suppressive microenvironment. Despite advances in surgery, radiation, and chemo-therapy, prognosis has changed little over the past decades, with a median survival of only 12–18 months (1,2).

Immunotherapies, particularly cellular approaches such as chimeric antigen receptor (CAR) T cells, natural killer (NK) cells, and tumor-infiltrating lymphocytes, have shown encouraging results in other cancers but face major challenges in GBM (3–5). These challenges include limited trafficking of immune cells across the blood–brain barrier (BBB), potent immunosuppression within the tumor microenvironment, and pronounced intratumoral heterogeneity (5). Overcoming these barriers requires innovative cellular platforms capable of entering the central nervous system (CNS) and engaging tumor tissue effectively.

Innate lymphoid cells (ILCs) represent a relatively recent addition to the immune landscape. Paralleling T cell subsets but lacking antigen specificity, ILCs are tis-sue-resident sentinels that contribute to immune surveillance, repair, and inflammation (6,7). Recent studies have highlighted ILCs as emerging players in cancer immunology, although their precise roles remain underexplored, particularly in GBM (8–11). Among these, group 2 innate lymphoid cells (ILC2s) are of special interest due to their roles in tissue repair, immune modulation, and cross-talk with both innate and adaptive immune compartments (12–14). However, whether ILC2s can infiltrate CNS tumors and persist within the glioblastoma microenvironment has not been experimentally demonstrated.

Emerging evidence that ILCs can migrate beyond mucosal and barrier tissues raises the possibility that ILC2s might be harnessed for brain-directed immunotherapy (10,15). Here, we report the first adoptive transfer of bone marrow–derived ILC2s into a mouse model of orthotopic GBM. Using CFSE labeling, we tracked their migration into the brain and tumor. Our findings suggest experimental proof that ILC2s can access and engraft within CNS tumors, establishing a foundation for developing ILC2-based cellular therapies. This work supports a new cellular platform that may be engineered or modulated for therapeutic purposes and lays the groundwork for future studies investigating the mechanistic and translational potential of ILC2s in glioblastoma.

## Materials and methods

### Animals and Ethics

Wild-type male C57BL/6 mice (10–12 weeks old; n = 6 from three independent co-horts; Jackson Laboratories, Bar Harbor, ME) were used. Animals were maintained under standard housing conditions with ad libitum access to food and water. All procedures were approved by the Augusta University Institutional Animal Care and Use Committee [IACUC #2011-0062] and conducted in accordance with the NIH Guide for the Care and Use of Laboratory Animals. The study was adhered to the ARRIVE guidelines for reporting animal research (ARRIVE 2.0 Essential 10 guidelines).

### Orthotopic GBM model

The luciferase-expressing GL261 line (GL261-Luc2) used in this study was obtained from the Georgia Cancer Center Cell Repository. This line is derived from the parental GL261 glioma cell line (NCI/ATCC origin) and stably transduced with firefly luciferase, allowing reliable in vivo bioluminescence imaging. GL261-Luc2 cells were cultured under standard conditions and stereotactically implanted into the right striatum (3 × 10_ cells in 3 µL PBS) of C57BL/6 mice as described previously (1). Mice were anesthetized using 3% isoflurane for induction and maintained with 1.5–2% isoflurane throughout surgical procedures. Tumor establishment and progression were verified by in vivo bioluminescence imaging (IVIS Spectrum). At the endpoint, animals were euthanized by CO_2_ inhalation followed by cervical dislocation, in accordance with the American Veterinary Medical Association (AVMA) Guidelines for the Euthanasia of Animals.

### Isolation and adoptive transfer of bone marrow-derived ILC2s

Bone marrow cells were harvested from donor C57BL/6 mice as described previously (16). Lineage-negative (Lin^−^) ILC2s were isolated using magnetic negative selection (BioLegend antibodies against CD3ε, CD5, CD19, LY6G, NK1.1, CD11b, CD11c, and Ter119) followed by Streptavidin MicroBeads (Miltenyi Biotec). Purified ILC2s were expanded for six days in RPMI-1640 medium supplemented with 10 ng/mL recombinant IL-7 and IL-33 (R&D Systems). ILC2 identity was confirmed by flow cytometry (Lin^−^CD45^+^Sca-1^+^ST2^+^GATA3^+^CD25^+^ICOS^+^c-Kit^+^IL-5^+^IL-13^+^). Prior to adoptive transfer, activated ILC2s were labeled with 5 µM carboxyfluorescein succinimidyl ester (CFSE; Invitrogen) for 10 minutes at 37°C, following established protocols (17-19). CFSE labeling provides a stable, heritable fluorescent marker that allows definitive tracking of trans-ferred cells in recipient animals without requiring re-staining for phenotypic markers [17-19]. To eliminate the possibility of free CFSE contributing to tissue fluorescence, la-beled ILC2s were washed three times with complete medium until the supernatant ex-hibited no measurable fluorescence. CFSE is an amine-reactive, intracellularly retained dye and does not dissociate once bound. Labeled ILC2s were intravenously injected via the tail vein (3 × 10□ cells/mouse) on day 9 post-tumor implantation, a time point ensuring established tumor growth. Control tumor-bearing mice received PBS injections. In flow cytometric analysis of recipient tissues, CD45^+^CFSE^+^ events represent transferred ILC2s, as CFSE was applied exclusively to the donor population. During downstream analyses, CFSE^+^ events were only considered ILC2s if they also expressed the expected immune phenotype (Lin^−^CD45^+^Sca-1^+^ST2^+^GATA3^+^). No CFSE^+^ events were detected in the brains of PBS-injected control mice, confirming signal specificity. This methodology is consistent with standard practice in adoptive immune cell transfer studies [17-19].

### Imaging and Flow cytometry

Bioluminescence imaging confirmed tumor growth, while fluorescence microscopy visualized CFSE^+^ ILC2s in the brain, meninges and peripheral organs (spleen, lung). Single-cell suspensions from these tissues were analyzed for CFSE^+^ ILC2s and other ILC subsets using a NovoCyte Quanteon flow cytometer (Agilent Technologies). Data were processed with FlowJo software (V10) as described previously [7]. Fluorescence quanti-fication was performed as mean pixel intensity (MFI) per microscope field. For each mouse, five non-overlapping regions were imaged at identical exposure settings, and the CFSE fluorescence channel was extracted and background-subtracted using ImageJ. The average pixel intensity per field per animal was used for statistical comparisons.

### Bioluminescence quantification

Tumor-associated photon emission was quantified using Living Image 4.7 software. For each mouse, a fixed-size region of interest (ROI) was drawn over the tumor, and an identical ROI was placed over a non-tumor brain region to obtain background signal. Background-subtracted radiance (photons/sec) was used for all comparisons. All imaging sessions were performed using identical exposure settings. Tumor-bearing mice included n = 6 per condition, derived from three independent experimental cohorts.

### Statistical analysis

All data were analyzed using GraphPad Prism (version 9). Frequencies of CFSE^+^ ILC2s in brain, meninges, spleen, and lung were compared using one-way ANOVA followed by Tukey’s post-hoc test. Quantification of CFSE fluorescence intensity in tissue sections was based on mean pixel intensity per field and analyzed using one-way ANOVA. For tumor bioluminescence, background-subtracted radiance values were compared between groups using unpaired t-tests, whereas longitudinal comparisons within the same animals used paired t-tests. All values are reported as mean ± SEM, and statistical significance was defined as p < 0.05.

## Results

### ILC2s traffic to and infiltrate brain tumors following adoptive transfer

Bioluminescence imaging confirmed reliable establishment of orthotopic GBM tumors by day 9 post-implantation, providing a stable model for subsequent ILC2 transfer (Figure 1A). At this stage, CFSE-labeled bone marrow–derived ILC2s were intravenously administered to tumor-bearing mice to assess their migratory behavior and CNS entry. At 96 hours post-transfer, fluorescence imaging revealed distinct CFSE^+^ signals within the meninges, brain, and tumor parenchyma (Figure 1B). Flow cytometric analysis confirmed that these infiltrating cells retained canonical ILC2 markers (Lin^−^CD45^+^Sca-1^+^ST2^+^GATA3^+^), identifying them as a discrete population within the glioblastoma microenvironment (Figure 1C). This represents, to our knowledge, the first direct indication that adoptively transferred ILC2s can cross the blood–brain barrier and localize within glioblastoma tissue following systemic delivery. To ensure that the CFSE^+^ signals represented intact ILC2s rather than free dye, we implemented multiple verification steps. CFSE binds covalently to intracellular amines and does not dissociate once incorporated; labeled ILC2s were therefore washed repeatedly until no detectable fluorescence remained in the supernatant. The CFSE^+^ events identified in brain and tumor tissue also co-expressed canonical ILC2 markers (Lin^−^CD45^+^Sca-1^+^ST2^+^GATA3^+^), a phenotype that cannot arise from free dye (Figure 1D). In tissue sections, CFSE^+^ structures displayed discrete, nucleated, cell-like morphology rather than diffuse extracellular staining, and no CFSE signal was observed in PBS-injected tumor-bearing controls. Together, these findings support that the CFSE^+^ signals reliably reflect transferred ILC2s.

**Figure 1.**
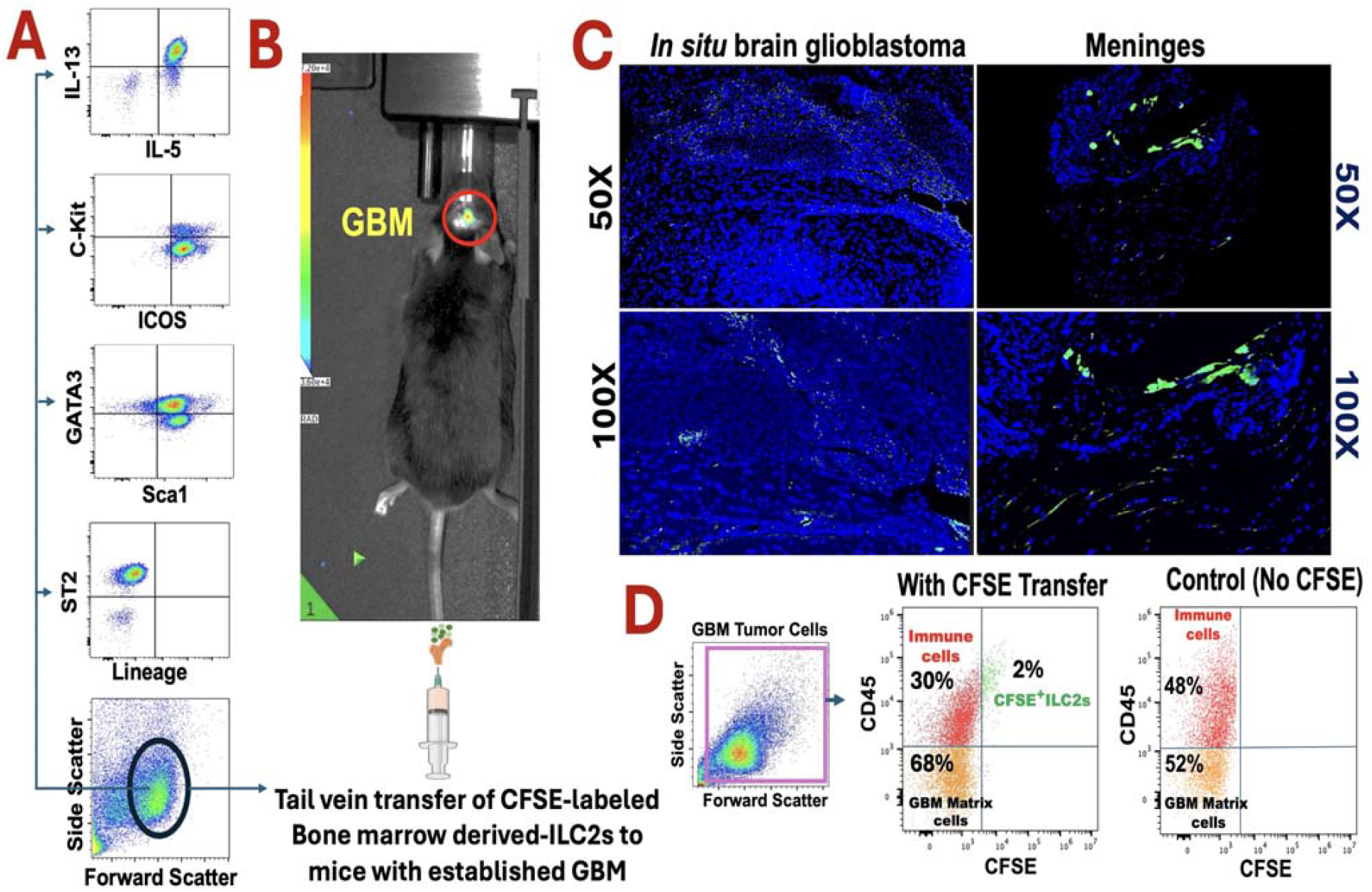
Adoptively transferred bone marrow–derived ILC2s retain canonical phenotype and infiltrate glioblastoma after systemic delivery. A) Phenotypic and functional characterization of bone marrow–derived ILC2s prior to CFSE labeling and transfer, showing expression of key ILC2 markers (IL-5^+^/IL-13^+^, ICOS^+^/c-Kit^+^, Sca-1^+^/GATA3^+^, ST2^+^ and lineage-negative profile). B) Experimental workflow and in vivo bioluminescence imaging confirming reliable establishment of orthotopic GL261-Luc glioblastoma prior to ILC2 adoptive transfer (day 9 post-implantation). CFSE-labeled ILC2s were injected intravenously via the tail vein. C) Representative fluorescence microscopy 96 hours after transfer showing CFSE+ ILC2s (green) within the glioblastoma parenchyma and meninges at 50× and 100× magnifications; nuclei are counterstained with DAPI (blue). D) CD45^+^CFSE^+^ cells represent the transferred ILC2 population. CFSE labeling was applied exclusively to donor ILC2s prior to transfer and serves as a definitive tracking marker for these cells in recipient animals. Pre-transfer characterization (Panel A) confirmed donor cell identity using canonical ILC2 markers (Lin^−^, IL-3^+^, c-Kit^−^, ICOS^+^, GATA3^+^, Sca-1^+^, ST2^+^). Control mice receiving no CFSE-labeled cells show no detectable CFSE signal, confirming specificity. This approach follows established methodology for adoptive cell transfer studies where unique cellular labeling provides unambiguous identification of transferred populations.

### Systemic and CNS-associated distribution of adoptively transferred ILC2s

Beyond the brain, CFSE^+^ ILC2s were also detected in peripheral tissues, including spleen and lung (Figure 2A), suggesting preserved systemic migratory potential. However, their frequency was markedly higher in the meninges and tumor regions (Figure 2B), suggesting a degree of preferential accumulation within CNS compartments. These findings indicate that ex vivo–expanded ILC2s maintain the capacity to traffic through the circulation and traverse the blood–brain barrier under the inflammatory conditions associated with GBM.

**Figure 2.**
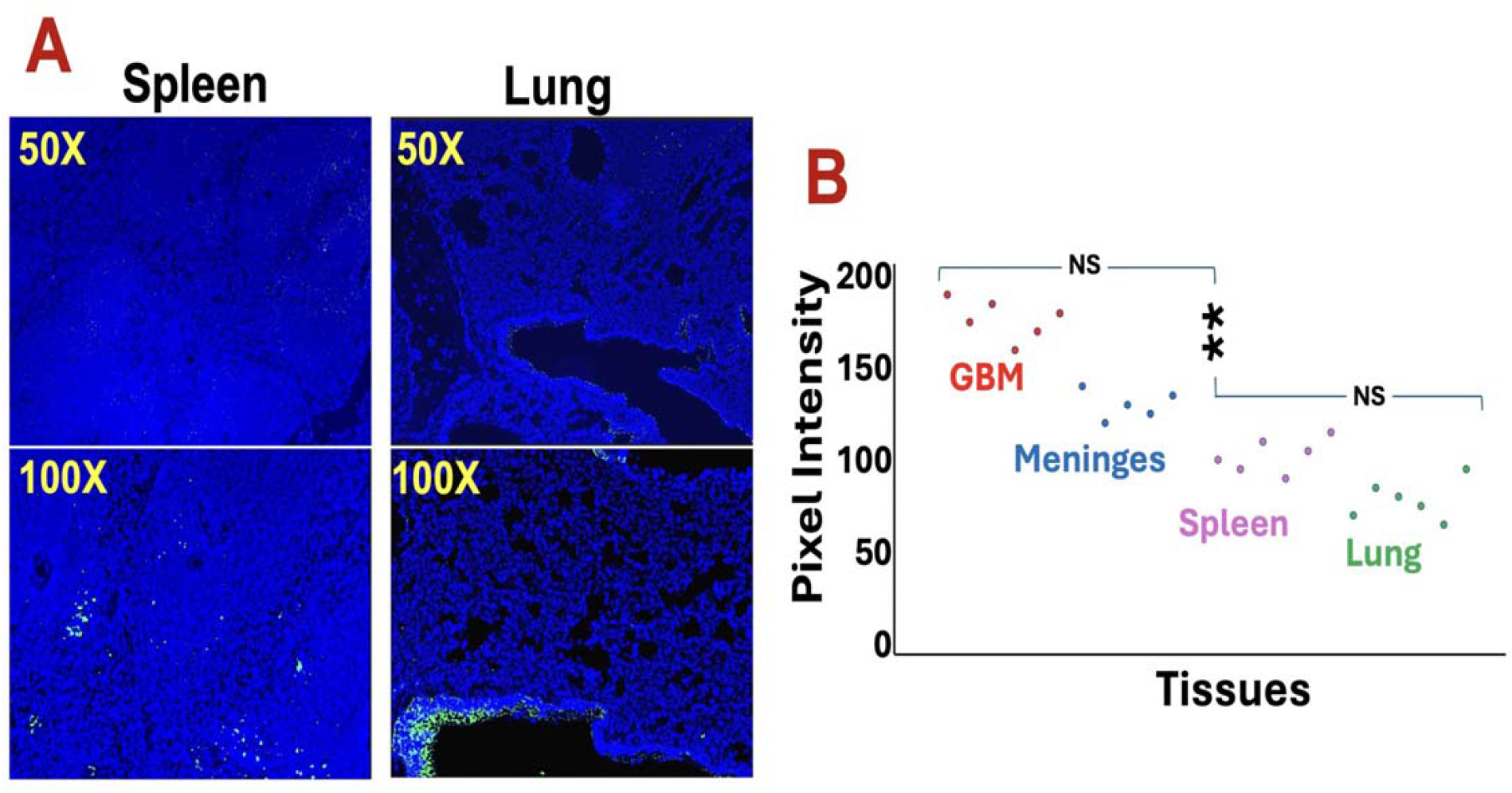

### Unmodified ILC2s infiltrate but do not alter glioblastoma growth

Despite successful homing and infiltration, ILC2 transfer did not significantly alter tumor progression as determined by longitudinal bioluminescence imaging (Figure 3A). Individual tumor radiance measurements at days 9 and 13 post-implantation showed comparable growth trajectories between ILC2-treated and control mice (Figure 3B), with no significant difference in tumor burden at either timepoint. These findings suggest that unmodified ILC2s alone are insufficient to exert antitumor activity within the GBM microenvironment. Nevertheless, their observed ability to enter and persist within intracranial tumors support a proof-of-concept for using ILC2s as a deliverable cell platform. Future efforts will focus on enhancing their therapeutic potential through cytokine priming, checkpoint modulation, or genetic engineering to improve tumor specificity and efficacy.

**Figure 3.**
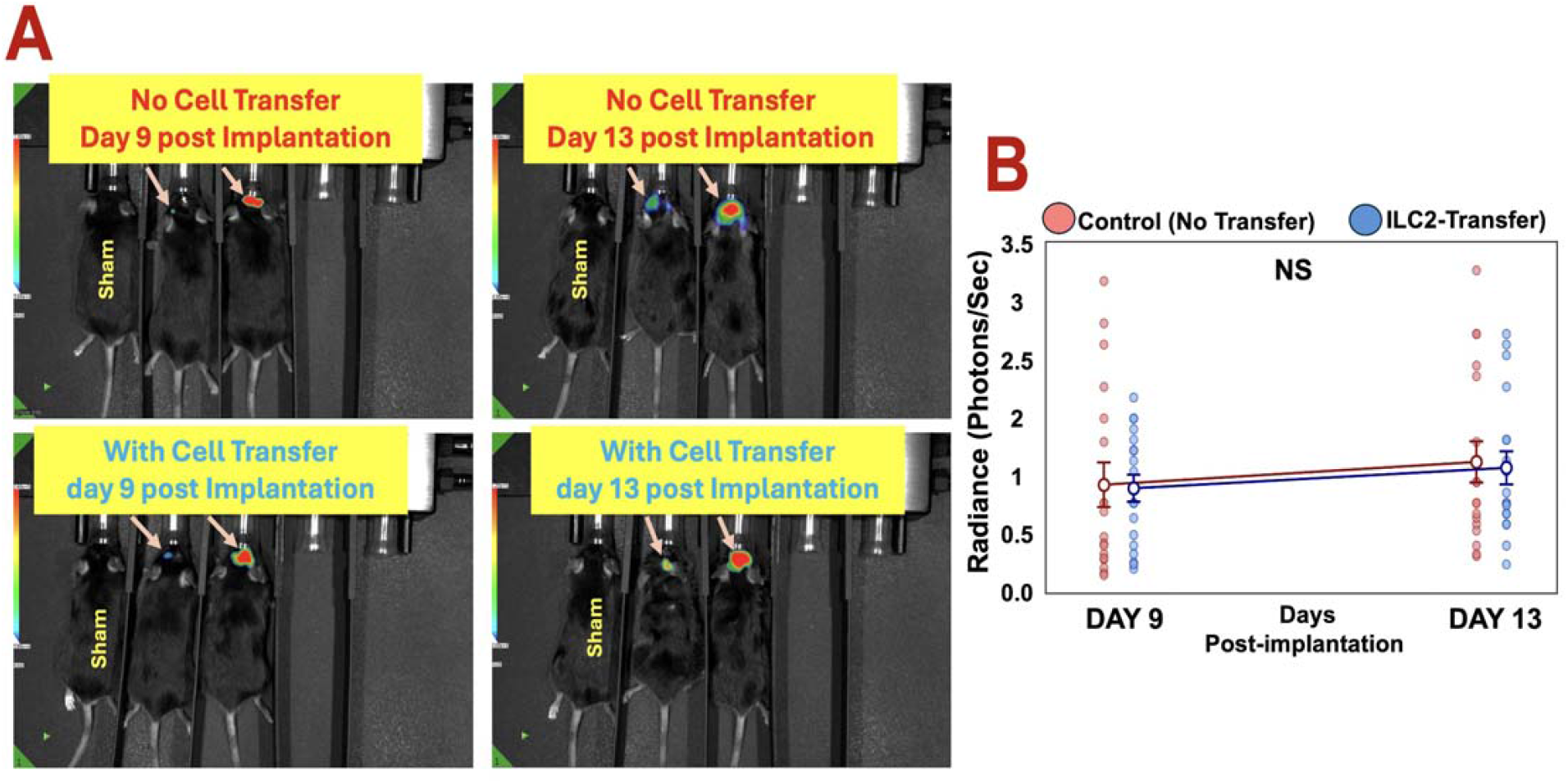
Unmodified ILC2s infiltrate but do not alter glioblastoma growth. A) Representative *in vivo* bioluminescence images of GL261-Luc tumor-bearing mice without ILC2 transfer (top) and with ILC2 transfer (bottom) on days 9 and 13 post-implantation. Sham-operated mice served as negative controls. Comparable bioluminescent patterns were observed between groups. (B) Longitudinal scatter plot showing individual tumor radiance values (photons/sec) for control (red) and ILC2-treated (blue) mice at days 9 and 13 post-implantation. Each filled circle represents an individual mouse measurement (n=18 per group across three independent experiments). Large open circles represent

## Discussion

This study provides the first experimental indication that bone marrow–derived ILC2s can traffic to and localize within glioblastoma tissue following systemic administration. While the adoptive transfer did not produce measurable differences in tumor size, the finding that ILC2s can successfully cross the blood–brain barrier and survive within the glioblastoma microenvironment represents an important conceptual advance for cellular immunotherapy of CNS tumors.

ILC2s have been increasingly recognized for their regenerative and immunomodulatory functions in peripheral tissues and solid tumors. Their ability to release type 2 cytokines, influence macrophage polarization, and promote tissue repair positions them as potential therapeutic agents to restore immune balance in the brain. Showing that systemically delivered ILC2s can access and persist within the glioblastoma niche provides a preliminary experimental foundation for future translational approaches.

As a proof-of-concept study, these experiments were limited by the short post-transfer observation period (96 hours) and the absence of pre-transfer cellular manipulations designed to enhance tumor specificity or effector function. This relatively brief window also limits our ability to determine how long the transferred ILC2s persist in the glioblastoma microenvironment or whether their early infiltration translates into sustained functional activity. Extended time-course studies will be needed to evaluate longer-term persistence, adaptation, and potential antitumor effects. Extending the time course and applying targeted modifications to ILC2s, such as cytokine priming or receptor engineering, may help reveal their full therapeutic potential against glioblastoma.

This work also provides a practical model for testing how ILC2s interact with the tumor microenvironment and for evaluating strategies to enhance their efficacy, such as combining adoptive transfer with agents that modulate IL-33 or TGF-β signaling. In conclusion, our findings support the feasibility of ILC2 brain homing and provide, for the first time, a proof-of-concept framework for developing ILC2-based adoptive cell therapies targeting glioblastoma and other CNS malignancies. Acknowledging the short observation period underscores that these results reflect an early stage of biological insight rather than a comprehensive therapeutic evaluation. Future studies will be essential to define the molecular cues guiding ILC2 migration, persistence, and functional programming within the brain tumor microenvironment.

## Conclusion

This brief report provides the first experimental evidence that systemically administered ILC2s can cross the blood–brain barrier and home to glioblastoma. Although the adoptive transfer of unmodified ILC2s did not alter tumor progression, their ability to infiltrate and persist within the brain suggests a new foundation for ILC2-based immunotherapeutic strategies targeting CNS malignancies. These findings open opportunities to explore engineered or cytokine-primed ILC2s as novel vehicles for brain-directed immunomodulation.

## Acknowledgements

Authors are thankful to Biomedical Research Associates of Georgia for providing help in the analysis, processing the data and optimizing the technical details.

## Statements & Declarations

### Declaration of AI Assistance

AI-based software (e.g., ChatGPT) was used solely for language refinement and grammar editing; all scientific content and interpretation were entirely developed by the authors.

### Author contributions

Conceptualization, L.P.W., B.B.; methodology, L.P.W., B.B., É.L.S., and J.C.Y.; software, B.I.B.; and S.E.N.; validation, B.B and L.P.W.; formal analysis, L.P.W., B.B., É.L.S., and S.E.N; investigation, B.B., N.Y. and L.P.W.; resources, B.B., L.P.W., and N.Y.; data curation, B.I.B., É.L.S., S.E.N. and A.A.; writing—original draft preparation, B.B., and L.P.W.; writing—review and editing, B.B., L.P.W., J.C.Y.; B.I.B., É.L.S., S.E.N., A.A., and N.Y., visualization, B.I.B., S.E.N., and B.B.; supervision, B.B., L.P.W. and É.L.S.; project administration, B.B., É.L.S. and S.E.N; funding acquisition, B.B., L.P.W. and N.Y.; All authors have read and agreed to the published version of the manuscript.

### Data availability

The original contributions presented in this study are included in the article. Further inquiries can be directed to the corresponding authors.

### Ethics approval

Animal study protocol was approved by the Institutional Animal Care and Use Committee (IACUC) of Augusta University (protocol code IACUC #2011-0062, Approved on 27^th^ of February 2023).

## Abbreviations

The following abbreviations are used in this manuscript:

GBM: Glioblastoma
ILCs: Innate Lymphoid Cells
CFSE: Carboxyfluorescein succinimidyl ester CNS Central Nervous System

